# Mapping climate risks across global pelagic fishing grounds

**DOI:** 10.1101/2025.05.07.652593

**Authors:** Matthew P. Faith, Angus Atkinson, Ryan F. Heneghan, Clare Ostle, Jose A. Fernandes-Salvador, Murray S. A. Thompson, Camila Serra-Pompei, Yuri Artioli, Katrin Schmidt, Sian Rees, Matthew Holland, Abigail McQuatters-Gollop

## Abstract

Ocean warming is projected to reduce the capacity of the global ocean to support fish biomass. The severity of projected biomass declines, however, is distributed highly unevenly at the global scale, meaning some nations face disproportionate socioeconomic risks from resultant fisheries declines. Here, we apply the IPCC climate risk framework through the lens of fishing grounds to identify global hotspots where climate-driven declines in supportable pelagic fish biomass pose the greatest threat to socioeconomic security. By integrating empirical size-spectrum projections of fish biomass declines (hazard) with present-day fishing effort (exposure) and a novel area-based index of fishing ground socioeconomic vulnerability, we identify risk hotspots around Southeast Asia, the western seaboard of Africa, and the West Pacific. Our projections are corroborated by an ensemble of ecosystem models and compared with historical observations of declining zooplankton biomass in the Northeast Atlantic, suggesting climate-driven fish biomass declines are already underway. Many of the regions we assessed as highest-risk are predominantly targeted by nations with relatively low adaptive capacity and who therefore face the greatest barriers to redistributing fishing effort to lower-risk fishing grounds. Socioeconomic risks could thus be partly mitigated by targeting risk hotspots with area-based interventions that directly support fish stocks, such as the restoration of essential fish habitats and the designation of Marine Protected Areas. Resourcing the development of regionally tailored models for high-risk regions will provide the best evidence for designing specific area-based measures that effectively reduce societal risks, but enforcing such measures will depend upon addressing illegal, unregulated and untracked fishing.

## 1 Introduction

Capture fisheries directly support the nutritional and economic security of billions of people [1]. Alongside being an important human protein source [2], fisheries provide essential micronutrients [3], including omega-3 fatty acids, and can provide feed for aquaculture and livestock [4, 5]. However, multiple human pressures threaten fish stocks, including overfishing, eutrophication, habitat destruction, and climate change [6–9]. Long-term ocean warming is a key climate effect impacting fish stocks, where rising sea surface temperatures and associated surface-nutrient depletion have already led to ocean-scale declines in phytoplankton biomass [10–12] – a trend which is set to continue with future climate change [13]. The food web impacts of decreasing phytoplankton biomass are compounded by shifts in the size structure of phytoplankton communities [14], reducing the efficiency of marine food webs and ultimately the supportable biomass of higher trophic levels, including fish [15]. While the complexities of disentangling the effects of climate change and fishing pressure make it challenging to attribute historical fish stock declines to climate change [16], multiple global projections agree that continued warming will cause a global decline in the supportable biomass of fish [15, 17–19].

To support policy decisions that increase climate-resilience, the Intergovernmental Panel on Climate Change (IPCC) periodically assess the potential adverse societal impacts arising from climate change as ‘climate risks’ [20]. Critically, risks are not only driven by the intensity, frequency, and distribution of climate hazards, but also patterns of societal exposure and vulnerability [21, 22]. In this context, the potential adverse societal impacts from climate-driven fish biomass declines can be considered a climate risk (fisheries climate risks), determined by patterns of societal exposure (the spatial overlap between society and fish biomass declines) and vulnerability (socioeconomic sensitivity to fisheries declines and regional adaptive capacity). The global distribution of fisheries climate risks is projected to be highly uneven [23, 24], with some areas of the ocean projected to see relatively more substantial declines in fish biomass [15, 17] and with socioeconomic dependence on fisheries varying substantially between countries [25, 26]. Some of the most fisheries-dependent nations, including some Small Island Developing States, are situated in regions projected to see relatively larger declines in supportable fish biomass and thus face disproportionate socioeconomic risks [18, 23].

Existing fisheries climate risk assessments have, to date, largely focussed on averaging projected biomass declines across individual Exclusive Economic Zones (EEZs; [18, 23]). These country-level projections support national-level climate adaptation strategies which aim to reduce societal vulnerability (e.g., livelihood diversification programmes; [27, 28]), alongside providing additional evidence to international climate mitigation negotiations of the societal harms which could be avoided from reducing greenhouse gas emissions. However, while emissions pathways are expected to determine the severity of fish biomass declines [17], targeted area-based measures, such as restoring essential fish habitats and avoiding overfishing (hereafter referred to as area-based fisheries mitigation measures), provide an additional opportunity to offset climate-driven fish declines and reduce societal risks [29–32]. Identifying global priority regions for targeting area-based fisheries mitigation measures requires an area-based approach to fisheries climate risk mapping, given the communities impacted by fisheries declines are not directly co-located with the areas of the ocean which provide fish. This is in part driven by the fact that fishing patterns are not spatially uniform across EEZs, but also because nearly a quarter (23%) of fish landings occur in Areas Beyond National Jurisdiction (ABNJ, or the High Seas) or in foreign territories [33]. Instead, mapping fisheries climate risks through the lens of fishing grounds could identify the specific ocean regions where fish biomass declines are projected to be substantial, and which are also most depended upon for the socioeconomic benefits provided by fisheries.

Here, we present a climate risk assessment of global pelagic fishing grounds. By combining Earth System Model outputs with a recently developed size spectrum-based approach [15] and existing ecosystem model ensembles (FishMIP; [17]), we project the ongoing declines in the supportable biomass of pelagic fish throughout this century (hazard). We couple these projections with present-day pelagic fishing patterns (exposure) and a novel index of the aggregate socioeconomic dependency and adaptive capacity of the nations operating within specific pelagic fishing grounds (vulnerability). By integrating these metrics through the tripartite IPCC risk framework, we create a pelagic fishing ground climate risk map that identifies where fish biomass declines pose the greatest threat to socioeconomic security, and which should be prioritised for area-based fisheries mitigation measures. Our risk mapping approach provides a framework for prioritising marine management efforts across the global ocean, including the High Seas, to safeguard food and economic security under climate change.

## 2 Results

### 2.1 Projected declines of pelagic fish biomass

Across much of the marine environment, ongoing ocean warming is projected to substantially decrease the supportable biomass of pelagic fish throughout this century (relative to a reference period of 1990-1999; Figures 1a-d). Modelled fish biomass decreases are driven by steepening size spectrum slopes (i.e., increasing dominance of small relative to large organisms; see Figure S1) coupled with decreasing phytoplankton biomass across much of the marine environment (see Figure S2). Even under a low emission scenario (SSP 1-2.6), notable declines of supportable fish biomass are projected by 2040-2049, with a global mean decrease of 8.5% (CI 6.3 – 10.5%; Figure 1a; see Figure S3 for mapped lower and upper confidence intervals). Under a high emission scenario (SSP 5-8.5), 2040-2049 declines in supportable fish biomass (Figure 1b) are projected to be similar to low emission declines, with a global mean decrease of 10.1% (CI 7.5 – 12.5%). By the end of the century (2090-2099) fish biomass declines are slightly dampened to a global mean decrease of 7.8% (CI 6.0 – 9.4%) under a low emissions scenario (Figure 1c). However, by the end of the century, a high emissions scenario is projected to result in mean average supportable biomass declines of 21.3% (CI 16.7 – 25.0%) with large areas facing substantially greater losses (Figure 1d).

**Figure 1.**
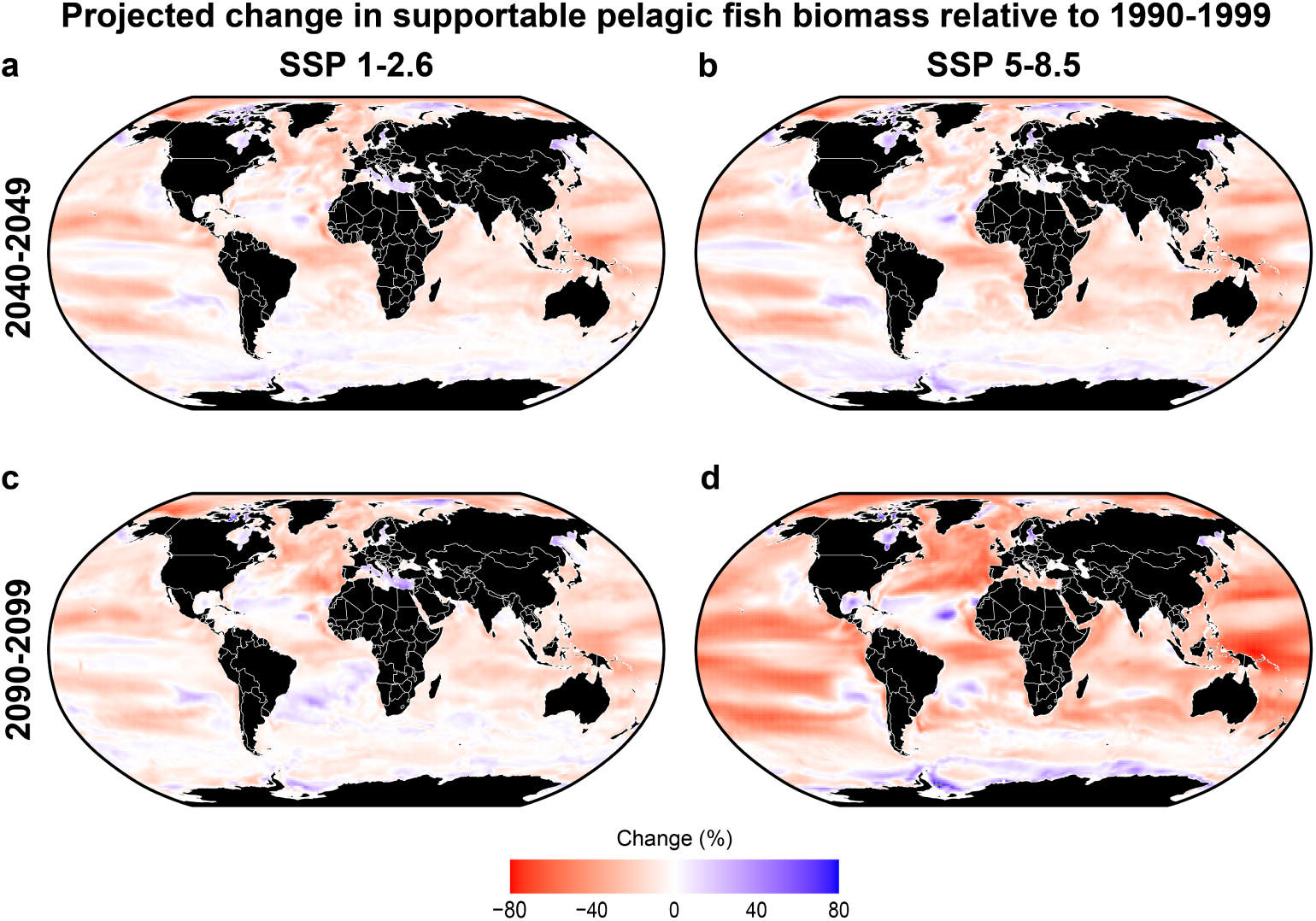
The effects of continued ocean warming on phytoplankton are projected to reduce the supportable biomass of pelagic fish. Percentage changes in total supportable pelagic fish biomass (g Cm^-2^) between 1990-1999 and 2040-2049 under a). SSP 1-2.6 and b). SSP 5-8.5; between 1990-1999 and 2090-2099 under c). SSP 1-2.6 and d). SSP 5-8.5.

### 2.2 Pelagic fishing ground risk assessment

We undertook a climate risk assessment of global pelagic fishing grounds by combining the projected trajectory of supportable pelagic fish biomass declines throughout this century (Figure 2a), with present day pelagic fishing patterns (Figure 2b), and the socioeconomic vulnerability of fishing activities within fishing grounds (Figure 2c). This identifies risk hotspots where climate-driven declines in supportable fish biomass pose the greatest threat to food and economic security. There is a great degree of overlap between areas modelled as undergoing large declines in fish biomass (Figure 2a) and those with intense pelagic fishing activity (Figure 2b), such as Southeast Asia, the Northeast Atlantic and the western coastline of South America. Small Island Developing States, such as the Federated States of Micronesia and Tuvalu, North Atlantic fishing nations, including Greenland and the Faroes, and several African nations, including Sierra Leone and Mozambique, were found to be among the most socioeconomically sensitive to fish biomass declines. Countries across Africa and the Middle East were then found to have the lowest adaptive capacity metrics, particularly Somalia, Yemen, and Eritrea. By combining sensitivity and adaptive capacity metrics into a single vulnerability indicator (see methods), we identified several African countries and Small Island Nations as the most vulnerable to climate-driven fisheries declines, given high socioeconomic dependence on fisheries (high sensitivity) and relatively less overall adaptive capacity (see Table S1 for full list of vulnerability indicators). These global inequalities translated into certain fishing grounds being disproportionately targeted by nations who are more vulnerable to fisheries declines (Figure 2c). Fishing grounds across Southeast Asia and the western Pacific, the eastern Atlantic, alongside hotspots in the Indian Ocean are targeted by the highest proportion of vulnerable nations.

**Figure 2.**
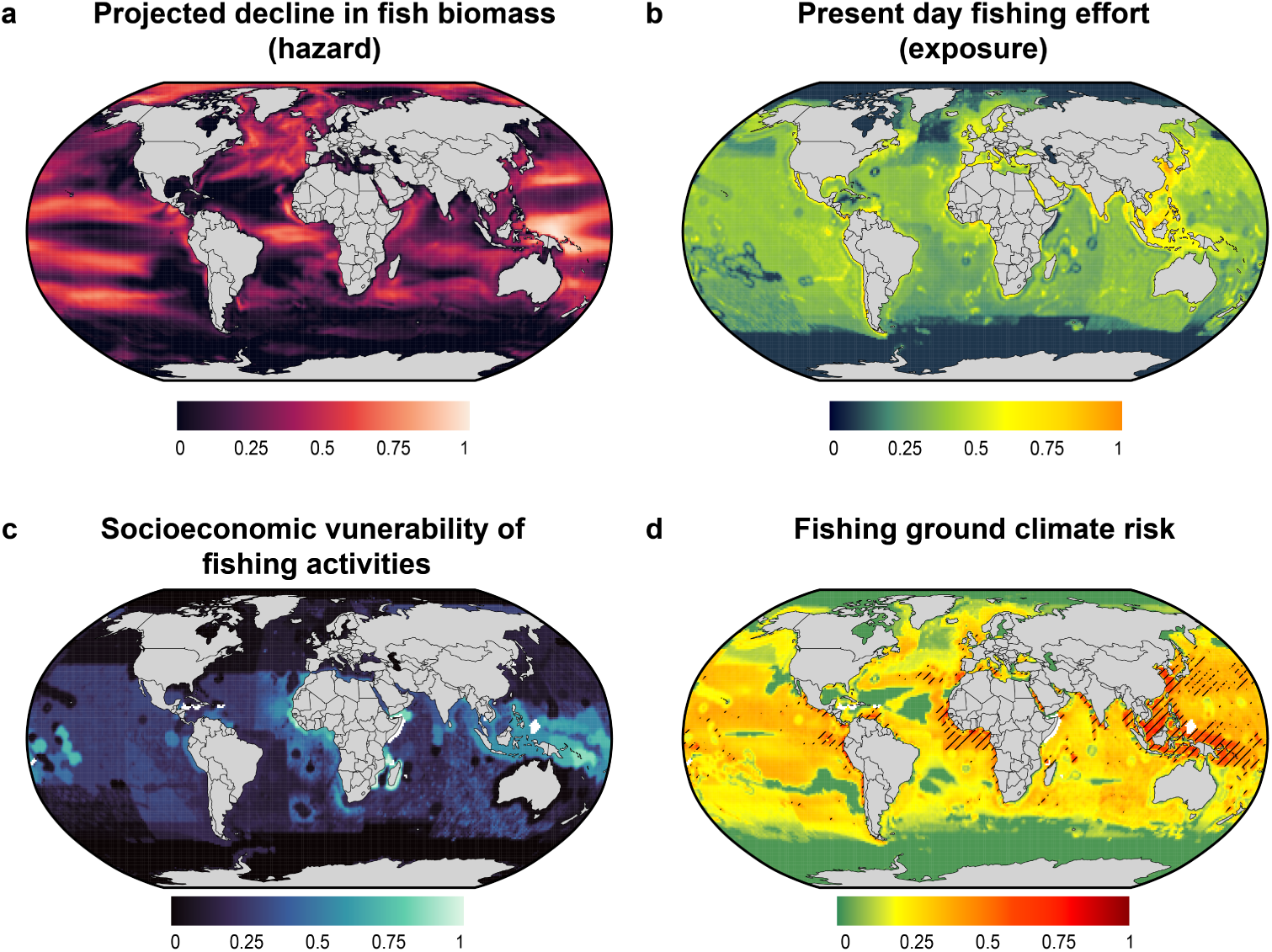
Pelagic fishing ground climate risk mapping, based on combining hazard, exposure and vulnerability, identifies priority regions for targeting area-based mitigation measures. A). Projected linear trend in supportable fish biomass decline between 1990-1999 and 2090-2099 under SSP 5-8.5 (hazard), derived from size spectra model; b). Mean total annual pelagic nominal fishing effort between 2014-2017 (exposure), processed from Rousseau *et al*. (2024). Exposure values were transformed (y = x^0.15^) to enhance visualization; c). Socioeconomic vulnerability of fishing activities (see methods); d). Overall fishing ground climate risk, calculated as the product of a, b, and c. Risk values were transformed to enhance visualization (y = x^0.15^) Hatching (d) indicates regions where both our size spectra projections and the ensemble mean of FishMIP projections (Tittensor *et al*., 2021) identified the location as high risk (90^th^ percentile). All metrics (a-d) were rescaled between 0 and 1 for plotting. Missing data are shown in white.

Our resultant fishing ground climate risk map (Figure 2d) identifies areas where modelled supportable pelagic fish biomass declines (Figure 2a) are co-located with intense pelagic fishing activity (Figure 2b), and which are targeted by nations with high socioeconomic vulnerability to fisheries declines (Figure 2c). With ongoing ocean warming, fish biomass declines across Southeast Asia and the western Pacific have the greatest potential to cause socioeconomic harm to the nations fishing there, and thus these fishing grounds were assessed as high risk. Other high-risk fishing grounds can be seen along the western coastline of Africa, and the western coastline of South and Central America. Medium-risk fishing grounds were identified with combined high-intensity fishing and substantial projected fish biomass declines but targeted by nations with lower overall vulnerability, such as the coasts of the UK and eastern coast of Canada, while other medium-risk regions are targeted by vulnerable fishing nations but are projected to see relatively moderate declines in fish biomass, such as the Maldives.

### 2.3 Validation of projections against ecosystem model ensembles and historical observations

To assess the robustness of our risk assessment, we compared results across a series of independent approaches. First, we compared our global projections of changing biomass of fish with the ensemble mean of mechanistic models from the Fisheries Model Intercomparison Project (FishMIP; [17]). The biomass projections are based on the relationship between plankton size spectrum slopes (normalised biomass size spectra; NBSS) and Chlorophyll-*a* (Chl-*a*) concentration [15]. We used this approach because pelagic size spectra are conceptually simple, well-grounded in ecological theory and observation, and the approach is empirical; it provides a method that is different to, and independent from, the suite of complex ecosystem models. Despite differences in the absolute magnitude of projected declines (Figure S4), these two independent approaches (NBSS and the FishMIP ensemble) agree on which fishing grounds are at highest risk (indicated by hatching in Figure 2d). Comparison between the FishMIP ensemble and NBSS approaches in the Northeast Atlantic further corroborates this agreement (Figure 3b, c, e). While the NBSS model projects a stronger decline than the ensemble mean, it falls within the uncertainty range of individual mechanistic models, and both approaches agree on the direction of significant declines (Figure 3e).

**Figure 3.**
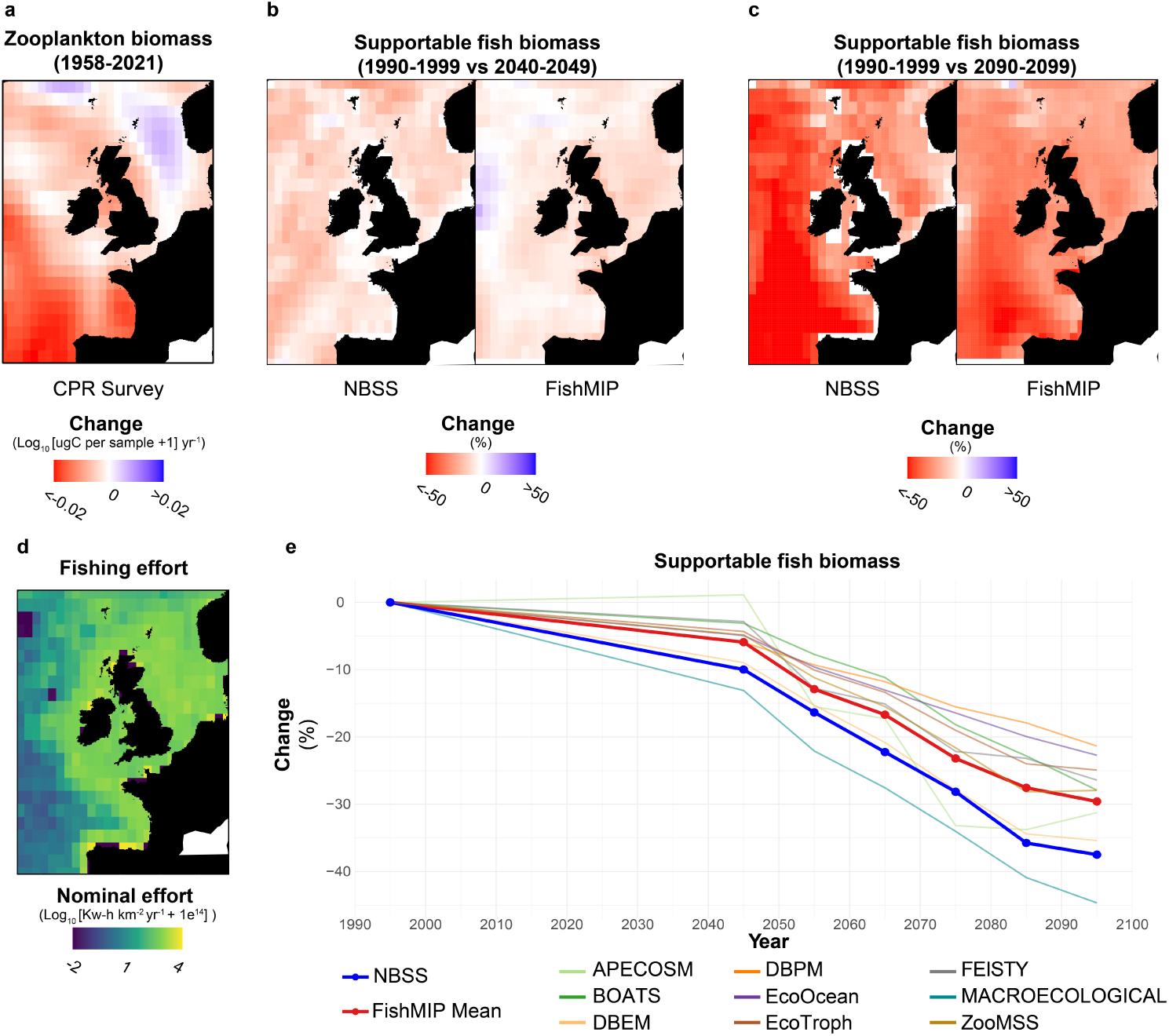
Size spectrum projections of declining fish biomass across the NE Atlantic are in-line with marine ecosystems model projections and are supported by the reductions in zooplankton biomass observed over the last 60 years of warming. a). Change in zooplankton biomass from observations over the last 60 years based on the Continuous Plankton Recorder (CPR) survey. These are compared on with projected changes in supportable fish biomass by b). 2040-2049 and c). by 2090-2099. Projections, are based on the ensumbles mean of ecosystem models from the FishMIP project (see method) and CMIP6 Earth system model output emission scenario (Shared Socioeconomic Pathway 5-8.5.). d) Mean total annual pelagic nominal fishing effort between 2014-2017, processed from Rousseau *et al*. e). Prohected chages in total values over time across the shole map domain for our method (NBSS) and the ensemble mean of FishMIP models, with individual FishMIP models marked (see Methods), Decadal averages were plotted as mid-decade (e.g., 1990-1999 is plotted as 1995).

The projections of changes in fish from both our size spectrum and the FishMIP ensemble are driven by projected changes in phytoplankton that is an output from Earth System Models (ESMs). However, global phytoplankton projections from ESMs also carry uncertainties [34, 35], and – importantly for our identification of risk hotspots – regional- and global-scale models can differ substantially [36]. We therefore compared the output of our NBSS-based projections of change in supportable biomass of fish (Figure 1) when driven both by the raw Chl-*a* and phytoplankton biomass outputs from ESMs and those from the European Regional Seas Ecosystem Model (NEMO-ERSEM; [37]). While as expected the ERSEM-derived output showed finer detail, the broader regionality of the trends in projected fish biomass decline was robust to the choice of the model used to derive the phytoplankton (see supplementary methods and Figure S5).

Importantly, our metric of declining fish biomass (Figure 3a) is a projected trajectory throughout this century (since 1990-1999), rather than a snapshot of the distant future. It essentially projects a continuation of the warming and loss of ecosystem efficiency that has already been observed over wide areas [14, 38]. If this projection that ongoing warming is reducing fish biomass is correct, historical warming should have already driven observable declines. Fish catch per unit effort has indeed decreased since the 1980s [39] but fish catch indices are often confounded by the complex interaction of climate and overfishing [16]. Zooplankton provide a clearer signal as arguably the closest unexploited trophic group to fish with sufficient data coverage. We therefore further validated our biomass projections at the regional scale in the Northeast Atlantic, a region selected due to substantial modelled fish biomass declines (Figure 1), intensive fishing activities (Figure 3d), and long-term zooplankton abundance time series data provided by the Continuous Plankton Recorder (CPR) Survey [40]. We calculated long-term trends in total zooplankton biomass across the Northeast Atlantic for the first time (for the period 1958-2021; Figure 3a). These trends indicate a general decline that is strongest in oceanic areas, directly overlapping with the regions of greatest projected loss in supportable fish biomass (Figures 3b, c). The observation that zooplankton, which consume phytoplankton and support fish, have declined during the last 60 years of warming [41] provides confidence that the supportable biomass of fish will continue to decline under future warming.

## 3 Discussion

Our projections of supportable fish biomass (Figure 1) reaffirm two key characteristics of climate-driven fish declines that underpin the need for effective fisheries climate risk management. First, while fish biomass declines are projected to be widespread, they are highly uneven at the global scale (Figure 1; [15, 17]), meaning some fishing nations face disproportionate socioeconomic risks [23]. Second, global supportable fish biomass is projected to decline even under strong climate mitigation scenarios (Figure 1; [17]). While the severity of future declines will largely depend upon international progress in reducing greenhouse gas emissions (Figure 1), the reality that significant warming is already locked into the global climate system means that some level of risks to fisheries will persist regardless of emissions pathways. Avoiding societal harms from climate-driven declines in supportable fish biomass will therefore depend upon multiple risk-reduction management avenues that address patterns of societal exposure and vulnerability, alongside identifying opportunities to reduce compounding pressures on fish stocks (Figure 4).

**Figure 4.**
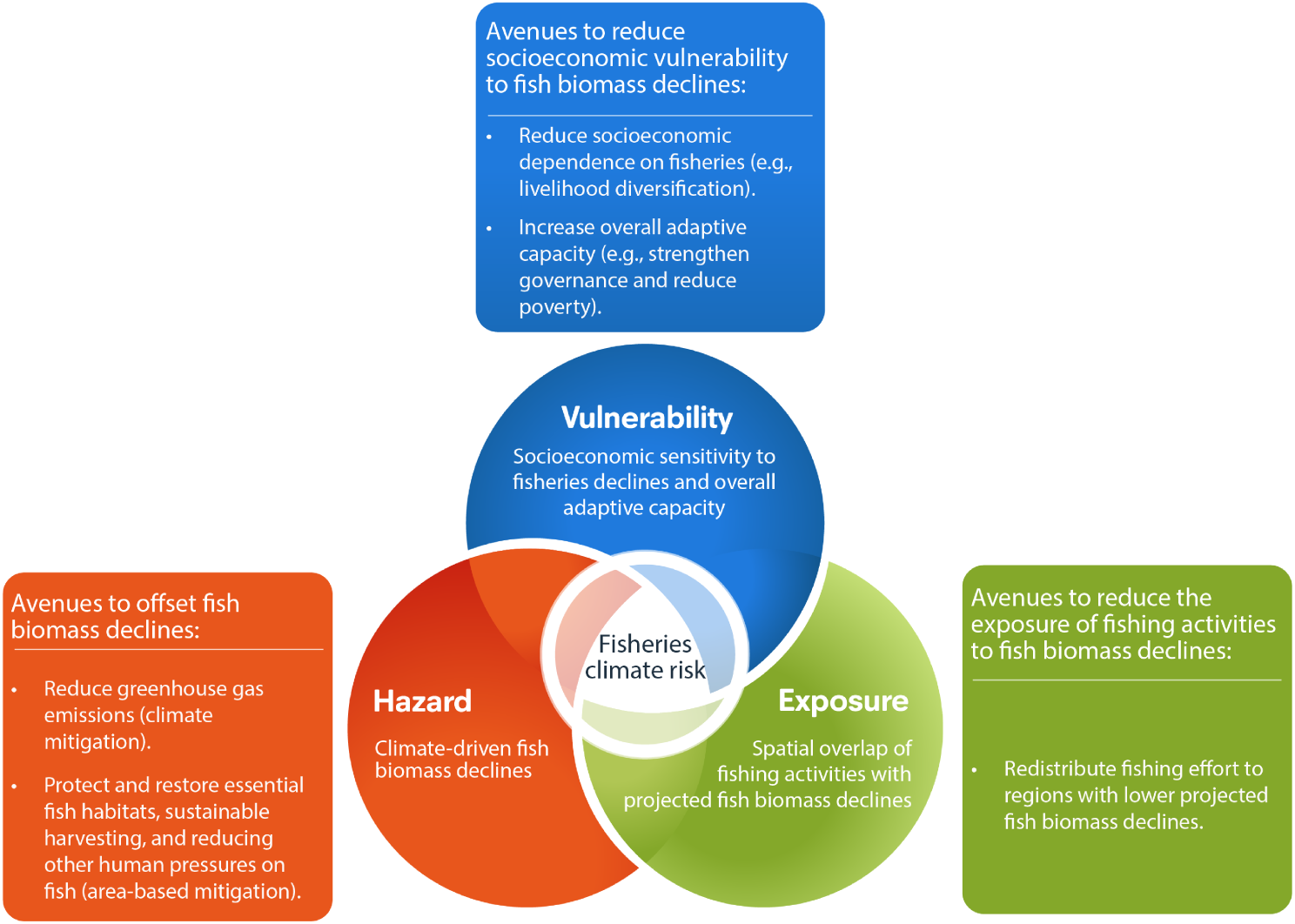
Applying the IPCC climate risk framework to declines in supportable fish biomass. Fisheries climate risk captures the potential for societal losses and damages arising from climate-driven declines in supportable fish biomass. The risk is mediated by the intersection of hazard (climate-driven declines in supportable fish biomass), exposure (present-day fishing patterns), and vulnerability (socioeconomic sensitivity to fisheries declines and overall adaptive capacity). Figure adapted from Figure TS.4 in the Technical Summary of the IPCC Special Report on the Ocean and Cryosphere in a Changing Climate (IPCC, 2019).

### 3.1 Multiple management avenues needed to reduce fisheries climate risks

Management measures that serve to reduce socioeconomic vulnerability to fisheries declines (Figure 4) can reduce the potential socioeconomic harms in nations that depend upon areas of the ocean we have assessed as high risk (Figure 2d). Efforts to reduce socioeconomic dependence on fisheries, such as through livelihood diversification programmes, alongside broader efforts to alleviate poverty and global socioeconomic inequalities (e.g., through the UN Sustainable Development Goals; [42]) are important approaches to reducing regional vulnerabilities to climate-driven fisheries declines [28, 43]. Considering projected fish biomass declines are highly uneven, and some regions are even projected to see increasing fish biomass (Figure 1), there may also be some opportunity to reduce exposure to fisheries declines through redistributing fishing effort to lower risk regions (Figure 4). Global fishing fleets have already been observed redistributing under the ongoing effects of climate change on fisheries [44], but the ability to redistribute fishing effort is highly dependent on regional adaptive capacity and is generally unfeasible for small scale fisheries [45]. Our vulnerability map (Figure 2c) considers both the socioeconomic sensitivity of individual countries to fisheries declines, but also their overall adaptive capacity. Global hotspots of fishing activity by vulnerable nations (Figure 2c), and thus regions with high overall fishing ground risk (Figure 2d), are therefore generally less likely to be able to avoid socioeconomic harms by simply shifting fishing effort to lower risk regions (Figure 2d).

Critical, therefore, is the fact that actual fish biomass declines are determined not just by climate-driven reductions in carrying capacity, but also by effective fisheries management and the wider conservation of marine ecosystems. While international progress in reducing greenhouse gas emissions is expected to determine the severity of carrying capacity declines (Figure 1; [17]), regional efforts to reduce compounding pressures on fish stocks and more broadly conserve marine ecosystems is an important tool for offsetting regional fisheries climate risks (Figure 4). By applying the IPCC climate risk framework to global fishing grounds (Figure 4), we identified global risk hotspots around Southeast Asia, the western seaboard of Africa, and the western Pacific (Figure 2d). Our approach identifies specific ocean regions where declining fish biomass intersects with intense fishing activities by countries with high socioeconomic vulnerability to fisheries declines, and so buffering fish stock declines in these regions has the greatest potential to offset adverse socioeconomic impacts.

### 3.2 A fishing grounds lens to marine ecosystem management under climate change

Although area-based marine management measures can only serve to buffer, rather than reverse, climate-driven declines in supportable fish biomass, such measures provide a vital tool for complementing other measures that reduce fisheries climate risks (Figure 4). Adopting sustainable practices in high risk regions (Figure 2), like balanced harvesting [46] and ensuring that overfishing is avoided, are particularly important given declining supportable fish biomass (Figures 1 and 3) is likely reducing the baseline levels of sustainable exploitation. Alongside responsive fisheries management, an increasing body of evidence demonstrates that wider marine conservations efforts can benefit fisheries under climate change. For example, actively restoring essential fish habitats can preserve complex age structures and increase recruitment [31, 32, 47], ultimately supporting fish stock climate resilience. Similarly, Marine Protected Areas (MPAs) can support fish stock climate resilience [29], and even directly benefit fisheries through spill-over effects [30].

Some of the fishing grounds we assessed as high-risk (Figure 2d) extend into Areas Beyond National Jurisdiction, highlighting opportunities for the newly adopted BBNJ Agreement [48] to establish conservation measures that co-benefit fisheries under climate change. The BBNJ Agreement serves to ensure the protection and sustainable use of marine biodiversity, providing a mechanism for designating biodiversity conservation measures, such as MPAs and other Area Based Management Tools (ABMTs; [48]). While the agreement was not established to directly manage fisheries, any resulting biodiversity-conservation measures have the capacity to co-benefit fisheries through spillover effects and increasing fish stock climate resilience [49, 50]. Our fishing ground risk maps (Figure 2d) therefore identify regions where any ABMTs designated under the BBNJ Agreement to conserve biodiversity could simultaneously mitigate fisheries climate risks. Some high-risk regions, such as Southeast Asia, are also known marine biodiversity hotspots [51], highlighting the potential for win-win area-based management measures that conserve biodiversity hotspots while buffering socioeconomic risks.

### 3.3 Addressing uncertainties for confident decision making

The validity of our risk maps is highly dependent upon the robustness of the underlying fish biomass projections. We used an empirical size-spectrum approach as a conceptually simpler and independent alternative to complex ecosystem models that carry notable uncertainties [52, 53]. Despite the radically different approaches there is a heartening degree of congruence: both our size spectra and the FishMIP ensemble identified similar risk hotspots (Figure 2d). A key uncertainty of our size spectrum approach is whether the same relationship between chlorophyll-*a* and the size spectra slopes, which underpins our projections [15], will hold under future environmental conditions. Atkinson et al. [15] used a diverse variety of natural aquatic ecosystems across the globe to determine the empirical relationship between size spectra slopes and chlorophyll-*a*, and it is important to note that these ecosystems were already facing a suite of climatic stressors and perhaps partially adapting to them in situ. This “natural laboratory” is a space-for-time substitution approach, but it provides some confidence that the relationship will not change with continued warming. Furthermore, our Northeast Atlantic case study provides further reassurance that, at least for this region, our projections are in line with historical observations of declining zooplankton biomass (Figure 3a), with the projections from FishMIP ensemble of mechanistic global ecosystem models (Figure 3b, c, e), and a more regionally tailored model (see Figure S5).

### 3.4 Barriers and opportunities to operationalising area-based mitigation measures

While our study identifies global priority regions for targeting area-based fisheries mitigation measures, the specific management measures likely to be most effective at the local-level will be highly context-dependent. Some fish stocks, for example, have spawning and catch areas that are not co-located, and so restoring essential fish habitats may provide fewer localised benefits than efforts to reduce overexploitation [54]. In addition, designating area-based management measures should not only consider declines in supportable fish biomass, but also the responses of specific stocks to climate change (e.g., through species redistribution; [55]). Some ecosystem models (e.g., SS-DBEM; [56, 57]) consider food web dynamics and thermal niches to project specific fish stock responses to ocean warming, yet producing reliable projections at finer scales depends upon inputs from regionally tailored biogeochemical models [57–59]. Developing regional models, however, is resource-intensive and so global coverage is limited [36]. We considered the overall adaptive capacity of countries fishing across global fishing grounds; some of the highest risk fishing grounds we identified in this study are also some of the most likely to face economic and technological barriers to developing regional models. By prioritising international funding and technological capacity building to support regional model development for the highest risk fishing grounds (Figure 2d), decisionmakers will be better equipped to implement area-based management measures that effectively offset fisheries climate risks.

The effectiveness of area-based fisheries mitigation measures will also depend upon addressing illegal, unreported, and unregulated fishing activities. Key risk hotspots we identified (Figure 2d), particularly across Southeast Asia and the eastern Atlantic, coincide with areas of intense untracked fishing [60]. High levels of illegal and unregulated activity are more likely to be associated with unsustainable levels of fishing [61], likely amplifying the potential societal harms from climate-driven fish stock declines within regions we assessed as high-risk. Addressing untracked fishing activities in high-risk areas is therefore a priority for avoiding the amplification of fisheries climate risks, but also to ensure that area-based management measures are enforced and effectively reduce societal risks.

## 4 Conclusions

Our fishing ground climate risk assessment demonstrates that while climate-driven declines in supportable fish biomass are globally widespread, the specific marine regions where these declines pose the greatest socioeconomic risk are highly concentrated. High risk hotspots are a result of substantial projected fish biomass declines in regions which are intensely fished by nations vulnerable to fisheries declines. The most vulnerable and highest risk regions identified in our study are generally depended upon by nations with relatively low adaptive capacity, and who therefore generally lack the technical and financial adaptive capacity to simply redistribute their fishing effort to lower risk regions. Significant global warming is already occurring and our analyses of the monitoring data available in Northeast Atlantic indicate that projected fish biomass declines are likely already underway. Marine conservation efforts, including restoring essential fish habitats and avoiding overfishing, must therefore be prioritised as a tool for proactively increasing fish stock climate resilience. Our novel fishing ground climate risk mapping approach identifies critical hotspots across South-east Asia, the western seaboard of Africa, and the western Pacific where targeted area-based mitigation measures have the greatest potential to offset societal risks. Risk hotspots extend into regions of the High Seas, underscoring the opportunities for the BBNJ Agreement to designate conservation measures that co-benefit fisheries under climate change. Developing regional ecosystem models tailored to the fine scale dynamics of regions we assessed as high risk will be critical to designing area-based mitigation measures that are locally effective. However, the effectiveness of mitigation measures will also depend on addressing illegal, untracked and unregulated fishing activities in high-risk fishing grounds that are likely amplifying societal climate risks.

## 5 Methods

### 5.1 Principle of using size spectrum slopes to estimate the supportable biomass of fish

Body size is an important trait in pelagic ecosystems, exerting control on key biological functions such as prey size and physiological rates [62, 63]. The slopes of Normalised Body Mass Size Spectra ([64]; hereafter “size spectrum slopes”) provide an index of the relative abundance of small and large pelagic organisms and therefore help to quantify the efficiency of mass transfer to large consumers [65]. Sprules and Munawar [66], Rossberg et al. [67] and Atkinson et al. [15] all found that systematic changes in size spectrum slopes across aquatic ecosystems related to their nutrient status. Atkinson et al. [15] further found that these size spectrum slopes did not relate to temperature as some have suggested, but only to the nutrient status of the systems, such that systems with low average Chl-*a* had steeply negative slopes (i.e. inefficient food webs) while those with higher Chl-*a* had less steep slopes and thus more efficient food webs. Importantly, these findings are all at the macroscale (i.e. of globally distributed in-situ ecosystems), with each presumably adapting to multiple stressors. For this reason, Atkinson et al. [15] used the relationship between size spectrum slope and annual mean Chl-*a* concentration in a space-for-time substitution approach, to estimate changes in size spectrum slope based on modelled changes in Chl-*a* derived by Earth System models. Since size spectrum slopes are broadly linear across the broad size spectrum of pelagic life [68], the size spectrum slope can be further used to estimate the supportable biomass of fish based on changes in modelled phytoplankton biomass [15].

Chl-*a* concentration (mg Chl-*a* m^-3^) and depth-integrated phytoplankton biomass (g C m^-2^) were obtained from GFDL-ESM4 and IPSL-CM6A-LR, two Earth System Models from the Coupled Model Intercomparison Project Phase 6 (CMIP6; [69]). These Earth System Models were selected because they are characterised by a very different Equilibrium Climate Sensitivity (ECS), with IPSL-CM6A-LR being one of the most sensitive (ECS = 4.6°C) and GFDL-ESM4 one of the least sensitive (ECS = 2.6°C) [70]. Both models provided monthly, global-scale data at 1° spatial resolution for the historical period 1970-2014, and for 2015-2099 under both low (SSP 1-2.6) and high emission scenarios (SSP 5-8.5). Chl-*a* and phytoplankton biomass values were averaged across the two Earth System Models, then averaged temporally for the periods 1990-1999 (see Figure S6 for historical baseline values) and decadal intervals between 2040 and 2099 (see Figures S2 and S7 for raw ESM projections).

Mean decadal Normalised Biomass Size Spectra slopes (*S*) were then calculated from projected mean decadal (Chl-*a*) values using the relationship derived in the meta-analysis of Atkinson et al. [15] (Equation 1). This empirical relationship was calculated from size spectrum slopes compiled from 41 distinct ecosystems ranging 600-fold in mean Chl-*a* values.

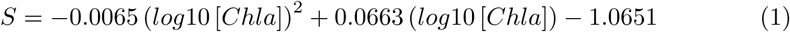

Following the approach of Atkinson et al. [15], our projected decadal averages of Normalised Biomass Size Spectra slopes (*S* values, Equation 1) and projected phytoplankton biomass values (mg C m^-2^) were used to project the total supportable fish biomass. This approach represents plankton as the size range 0.5 to 50,000 pg C and fish 0.5 g C to 50,000 g C [68]. To estimate the relative changes in biomass of fish size range compared to the phytoplankton size range, projected size spectra slope (*S*) is used to extrapolate projected phytoplankton biomass. Equation 2 summarises this approach, where the size range of fish is represented as 12 orders of magnitude greater than the phytoplankton size range. For each grid cell, the percentage change in supportable fish biomass was calculated for each future decadal interval (between 2040 and 2100) relative to the 1990 - 1999 baseline.

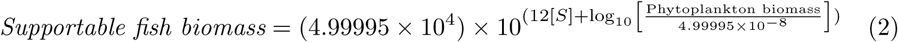

### 5.2 Risk assessment of fishing grounds

#### 5.2.1 Hazard metric: rate of declines of fish biomass

Decadal projected percentage changes in supportable fish biomass (relative to 1990-1999, see method above) were converted to rates of change in supportable fish biomass, using a least-squares linear regression (forced through the origin) for each grid-cell. Rates of change were filtered to only include regions of projected declining fish biomass, converted to positive values (multiplied by −1), and normalised between 0 and 1 using Equation 3 to produce Figure 2a. Our metric normalisation approach meant that we assessed relative, rather than absolute, hazard (and risk). We therefore did not compare risks under different emissions pathways but only included SSP 5-8.5 due to similar spatial distributions of relative fish biomass declines under each emission scenario (see Figure S8 for risk comparison between SSP 1-2.6 and 5-8.5).

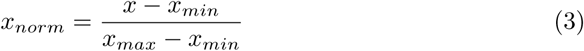

#### 5.2.2 Exposure metric: present day fishing effort

Patterns of present day pelagic fishing effort were sourced from the database provided in Rousseau et al. [71]. While other fishing effort datasets are freely available, such as from automatic identification systems (AIS) [72], the database from Rousseau et al. [71] provides data in regions with poor AIS data coverage, such as Southeast Asia [60], making it more appropriate for our global approach. Total annual average country-aggregated nominal fishing effort (kW-h km^-2^ y^-1^) was processed for the most recent four years available (2014 – 2017). While Rousseau et al. [71] provide data for the effective fishing effort (nominal effort corrected for relative developments in catch efficiency between countries), we opted to use nominal fishing effort due to its comparability to other available fishing databases, including AIS databases. The country-aggregated mean total annual fishing effort was normalised to a score between 0 and 1 for each grid cell using Equation 3.

Importantly, fishing effort was filtered to only include activities targeting pelagic functional groups given both our NBSS approach and the ensemble of ecosystem models used in FishMIP are most representative of pelagic food webs. This is partly due to our NBSS approach and several individual ecosystem models being primarily driven by surface plankton dynamics, but also because the finer scale processes that regulate coastal fisheries are poorly constrained in global models [17]. While our risk mapping approach is highly transferable to other fisheries (e.g., benthic), we only linked our biomass projections to pelagic fish functional groups to avoid introducing extra uncertainty into our analyses. Our exposure, vulnerability, and risk maps therefore inherently underestimate risks in regions dominated by demersal or benthic fisheries (see Figure S9 for exposure, vulnerability and risk maps based on all fishing activities).

#### 5.2.3 Vulnerability metric: socioeconomic dependence on fishing grounds

A composite metric of national vulnerability was calculated based on the IPCC risk framework [21], which considers vulnerability as a function of sensitivity and adaptive capacity. We therefore calculated Vulnerability (*V*) as:

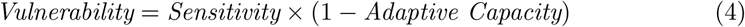

A sensitivity metric for each country was calculated according to the fisheries dependency metric presented in Blanchard et al. [73] and Barange et al. [25]. For each country, percentile ranks were calculated for the economic contributions of total marine fish landings, the employment contributions of all fisheries, and the food security provided by fisheries. A mean average score was then calculated from these equally weighted metrics before calculating a normalised score for each country using Equation 3. Percentage total GDP contributions from all fish landings were calculated using the mean total annual fish landings between 2015-2019 from the SeaAroundUs database [74] and the 2015-2019 average GDP values from the World Bank [75]. Eritrea and Venezuela did not have GDP data available for 2015-2019, so we used the most recently available GDP values (for the year 2011 for Eritrea and 2014 for Venezuela). Percentage workforce employment in the fisheries sector by country was calculated from the employment in fishing and agriculture indicator from the Food and Agriculture Organisation’s FAOSTAT database [76] and total working population (ages 15-64) values from the International Labour Organization’s ILOSTAT database [77], using the most recent year of data available for each value and country. Fisheries contributions to food security were calculated from Equation 5, as presented in Blanchard et al. [73] and Barange et al. [25], using updated country dietary profile data provided in Nash et al. [78]. A standard required animal protein intake value of 36 grams per capita per day was used [79], keeping our approach in-line with previous studies [25, 73].

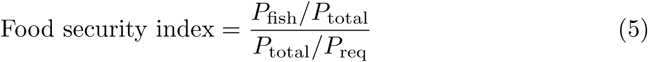

Where *P*_fish_ is the per capita fish protein intake, *P*_total_ is the total animal protein intake, and *P*_req_ is the required animal protein intake.

An Adaptive Capacity (AC) index was then used to address the broader ability of a nation to cope with fisheries declines through institutional stability and economic resilience. In a similar way to sensitivity, AC was calculated using three national metrics: GDP per capita (current US$; [80]); the Human Development Index (HDI; [81]); and the mean average of the six Worldwide Governance Indicators (WGI; [82]). To calculate AC, the most recent year of available data was extracted for each country across the three datasets. The HDI, GDP, and WGI values were then converted to percent ranks before calculating a mean average by country which was then normalised using Equation 3. While our sensitivity metrics are specific to the fisheries sector, we utilised broad national metrics for adaptive capacity. This reflects the premise that a nation’s ability to cope with sector-specific losses (i.e., fisheries declines) depends fundamentally on its broader institutional stability, economic wealth, and human capital to facilitate alternative livelihoods and food sources [25, 26, 83]. The final national Vulnerability (*V*) score was calculated using Equation 4, where higher AC scores reduce the overall vulnerability of a nation, followed by a final normalisation step using Equation 3.

Due to data availability issues, we calculated sensitivity and adaptive capacity indices for countries which did not have all three of the respective metrics available (by taking a mean average of the available indices), before renormalising between 0 and 1. 206 countries had data available for at least one of the individual metrics for both sensitivity and AC, whereas 106 countries had data available for all individual vulnerability metrics. This approach is likely to skew the final vulnerability score for countries with metrics missing (the six individual vulnerability metrics are not necessarily strongly auto-correlated) but provides a precautionary assessment solution [84], given the data available. Data could not be sourced for dietary, economic and employment metrics specific to pelagic marine fisheries so we used data for all fisheries (e.g., demersal and in some cases freshwater). Our sensitivity metrics therefore likely overestimate socioeconomic vulnerability to pelagic fisheries declines in countries that depend upon non-pelagic fish groups. We therefore calculated the final grid-cell vulnerability as a function of each country’s specific pelagic fishing intensity (see below), to scale down the influence of broad national fisheries vulnerability scores in areas where pelagic effort is low. In addition, because our vulnerability metrics rely on national-level reporting, they do not resolve socioeconomic inequalities within individual nations. This is particularly relevant to Overseas Territories that may be relatively more vulnerable to fisheries declines than their respective administering powers, yet aggregated reporting of socioeconomic datasets means these vulnerabilities cannot be robustly mapped [85–87].

To link national vulnerability to spatial patterns of fishing effort, we calculated the socioeconomic vulnerability of fishing activity for each grid cell. Country-level values of mean total annual (2014-2017) pelagic fishing intensity (days) per grid-cell were sourced from Rousseau et al. [71]. For each grid-cell, country-level pelagic fishing effort was multiplied by the country’s corresponding national vulnerability metric. These values were summed per grid-cell and divided by the total fishing effort in that cell to capture the effort-weighted socioeconomic vulnerability of fishing activities in each grid-cell (Equation 6).

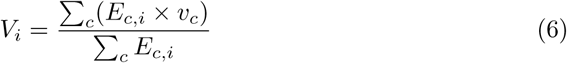

Where *V_i_* is the effort-weighted socioeconomic vulnerability of fishing activities in grid cell *i*, *E_c,i_* is the total pelagic fishing effort by country *c* in cell *i*, and *v_c_*is the national vulnerability metric for country *c*.

Limitations in data availability resulted in some countries with available fishing data in Rousseau et al. [71] missing vulnerability metrics. To represent fishing grounds with poor data coverage, grid-cells where the proportion of total pelagic fishing effort associated with vulnerability data was less than 50% were marked as missing data in our figures. In addition, the EEZs of countries with missing fishing data in Rousseau et al. [71] were also marked as missing data.

#### 5.2.4 Risk mapping

A fishing ground climate risk indicator was calculated for each grid cell as the product of the hazard, exposure and vulnerability metrics detailed above (Equation 7), based on definitions of risk from the IPCC [88].

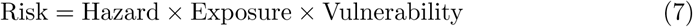

All underlying risk calculations were performed on linear data normalised between 0 and 1 (Equation 3). However, to account for the distinct statistical distributions of the individual risk metrics (Equation 7), data scaling was optimised for visualisation in Figure 2. Fishing effort (exposure) and thus overall risk are strongly skewed; these metrics were therefore transformed using a power of 0.15 (Figure 2b, d) to enhance visual contrast in lower-vulnerability regions. Hazard (biomass decline; Figure 2a) and vulnerability (Figure 2c) are presented on a linear scale.

### 5.3 Inter-model comparison

To assess the robustness of our fish biomass projections for downstream risk assessments, we compared our NBSS-derived risk estimates with risk scores derived from the ensemble mean of total consumer biomass (TCB) from the FishMIP models [17] under SSP 5-8.5. FishMIP simulations were driven by the same two Earth System Models used to make our NBSS projections (GFDL-ESM4 and IPSL-CM6A-LR) [17]. To ensure direct comparability, FishMIP outputs were processed using the identical hazard methodology applied to our NBSS projections. Decadal projected percentage changes were converted to rates of change using linear regression (forced through the origin), filtered to only include trends of declining TCB, and normalised between 0 and 1. These FishMIP-derived trend maps of TCB were then combined with the same exposure and vulnerability layers to generate comparable risk maps. We defined high-risk areas as grid cells falling within the upper decile (*≥* 90th percentile) of the global risk distribution for each respective dataset. This relative threshold approach accounts for differences in absolute magnitude between modelling approaches while preserving spatial ranking. The consensus layer (Figure 2d, hatching) identifies the spatial intersection of these hotspots, where both the NBSS model and the independent FishMIP ensemble concurrently classify the area as high risk.

### 5.4 Regional validation in the Northeast Atlantic

To provide a context and sense-check for the projections of changes in supportable biomass of fish we examine past changes in a well sampled region, the NE Atlantic, over the last 65 years which included phases of rapid warming [41, 89]. There were no total fish biomass data available over sufficiently large temporal and spatial scales, and the nearest trophic levels to fish with a suitable high-quality dataset were zooplankton from the Continuous Plankton Recorder (CPR) Survey (www.cprsurvey.org). Here, and for the first time, we have estimated long-term trends in total zooplankton biomass over the large areas covered by this sampling instrument.

To do this, zooplankton abundance counts per sample (i.e. no. per 3m^3^) between 1958 – 2021 were provided by the Continuous Plankton Recorder survey (www.cprsurvey.org, dataset: https://doi.org/10.17031/67ebed65bd6f9). The abundance data were multiplied by their respective carbon mass estimations provided in a new database [90] and then summed to obtain total biomass (µg per 3m^3^). The data were log transformed using log_10_ (µg per 3m^3^ + 1) in order to homogenise the variance and reduce the influence of zero counts [91]. These biomass data were then gridded onto a 1° x 1° grid, using the monthly mean value for each grid cell, and only using grid cells with *≥* 3 samples to reduce the potential impact of low sampling effort [91]. To visualise the changes within the Northeast Atlantic, the data were interpolated using an objective mapping technique, described by Equation 8.

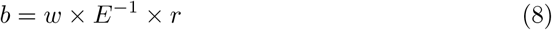

Where *b* is the mapped property, *w* is the data weights, *E* is the covariance matrix and *r* is the residuals (weighted mean) [92].

To further validate our projections in this region, we compared our NBSS-derived estimates fish biomass declines against estimate of total consumer biomass (TCB) declines from the FishMIP ensemble. We extracted spatially averaged decadal projections of TCB from the FishMIP database under SSP 5-8.5 for a region of the Northeast Atlantic (longitude −18° to 8.75°, latitude 41.5° to 63.75°). To create a continuous time series, projections periods were converted to mid-decade values (e.g., the 2050-2059 period was converted to 2055). We then calculated the spatially averaged percentage change in supportable biomass relative to the 1990–1999 baseline (i.e., 0% change at 1995) for both our NBSS projections and each individual FishMIP ecosystem model. This allowed us to compare the magnitude and rate of our projected declines against the inter-model uncertainty of established mechanistic approaches (Figure 3e).

## Supplementary information

Supplementary figures and country-level fisheries vulnerability metrics can be found in the supplementary file accompanying this manuscript.

## Acknowledgements.

MF acknowledges funding from the UK Department for Environment, Food and Rural Affairs (DEFRA), the University of Plymouth, and The Alan Turing Institute Enrichment Scheme. AMG, CO and MH acknowledge funding through the marine arm of the DEFRA Natural Capital and Ecosystem Assessment (NCEA) programme (NC34 Pelagic program - “PelCap”). AA was also funded through the European Space Agency ESA Cluster Multistressors Impact on Ocean Health grant number 4000142100/I-DT project Multiple Threats on Ocean Health (MiTHO), the NERC AtlantiS programme, and Defra PIECe programmes. JAFS contribution has been funded through the H2020 project OBAMA-NEXT (#101081642), as well as BioBoost+ project within the Biodiversa+ European Biodiversity Partnership program. Funded by the European Union. YA acknowledges funding from European Union’s Horizon 2020 research and innovation programme under grant agreement no. 820989 (project COMFORT), and from the NERC projects RECI-CLE (NE/M004120/1) and FOCUS (NE/X006271/1). MSAT was supported via the Natural Environment Research Council and the Economic and Social Research Council grant NE/V017039/1. RFH was supported by the Australian Research Council Discovery Early Career Researcher Award DE250100013. Current funding that supports the collection of CPR data in the North Atlantic includes: the UK Natural Environment Research Council: Atlantic Climate and Environment Strategic Science (AtlantiS) NE/Y005589/1, Department for Environment Food and Rural Affairs UK ECM-64770, National Science Foundation USA A101666/83643000, NERACOOS N21A01303, Fisheries and Oceans Canada F521A-210734/001/HAL, Horizon 2020: 862428 Atlantic Mission and AtlantECO 862923, IMR Norway and Ireland Marine Institute SERV-22-FEAS-090. Views and opinions expressed are however those of the author(s) only and do not necessarily reflect those of the funders.

## Data availability statement

All code used to generate the figures in this paper can be found at: https://github.com/matthewpfaith/FishingGroundRisk. Earth system model outputs can be found at: https://data.isimip.org/. For all other data, including socioeconomic and fishing effort data, please see the methods section.

## Author contributions

Conceived and designed the study: MPF and AA. Analysed the data: MPF, CO, and AA. Wrote the paper: MPF, AA, RH, CO, JAFS, CSP, YA, KS, SR, MT, MH, and AMG.

