## Supplementary figures and images for "Mapping climate risks across global pelagic fishing grounds"

### Figure S4

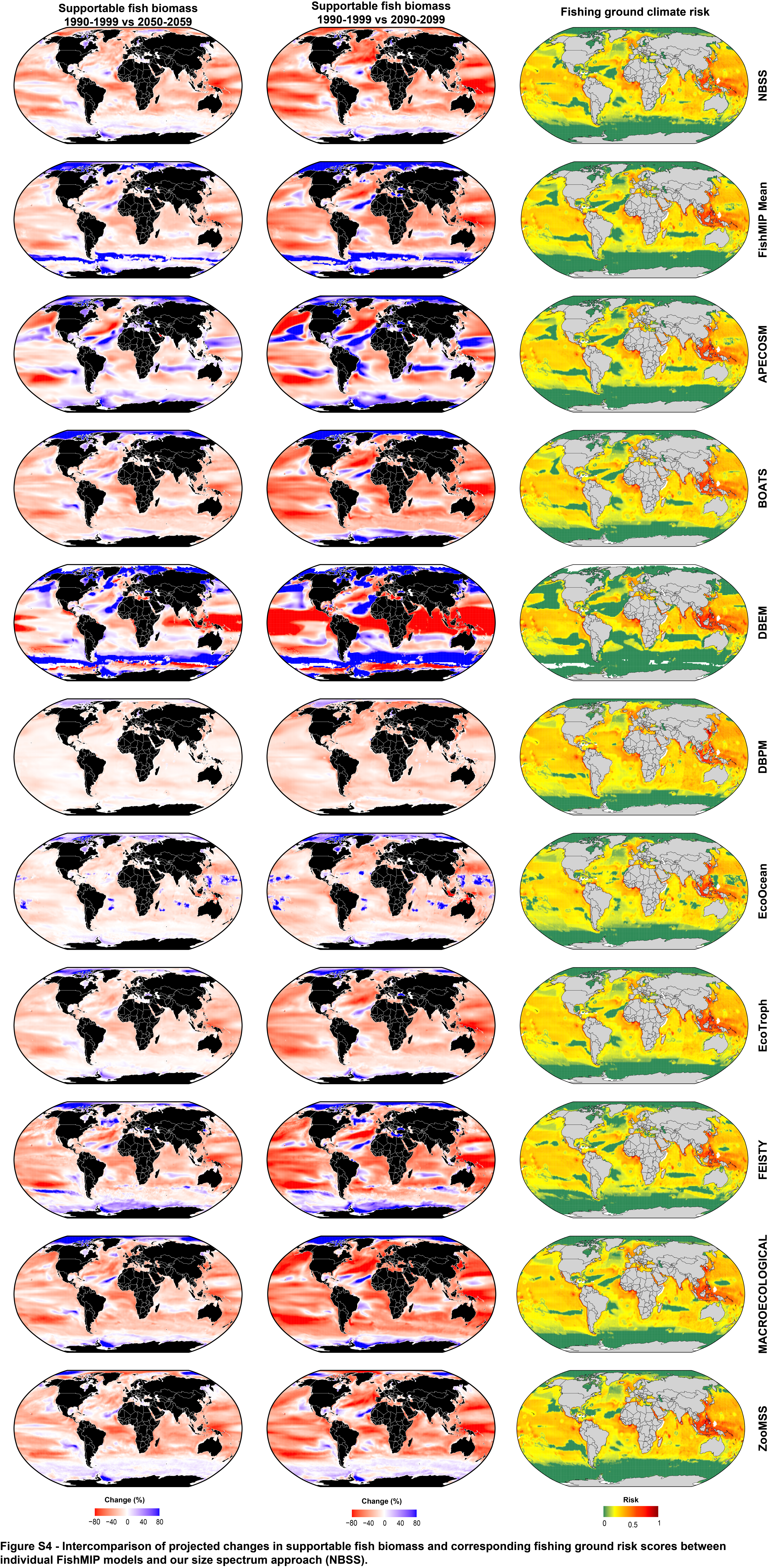
