## Supplementary Information for "Mapping climate risks across global pelagic fishing grounds"

### 1 Supplementary methods

#### 1.1 ERSEM comparison

Changes in phytoplankton used as inputs to our size spectrum-based projections are outputs from Earth System Models, which are known to have issues in localised shelf environments [1]. Therefore, to provide a further test of our projections, we compared outputs based on the NEMO-ERSEM modelling system [2] to the Earth System Models used in CMIP6, which were used for our global projections (Figures 1-3 in the main manuscript). Specifically, we used a downscaled projection driven by the Earth System Model IPSL-CM5-MR under the RCP8.5 scenario using the NEMO-ERSEM coupled model system on the Atlantic Margin Model at 7km resolution (AMM7) [3].

NEMO-ERSEM projections are made over much finer scales in comparison with CMIP6 models and include a greater range of physical and biological processes. NEMO [4] is a hydrodynamic model and ERSEM is an intermediate complexity ecosystem model with 3 zooplankton and 4 phytoplankton groups [2]. Outputs from the original 7km grid were aggregated on the common 1-degree grid before analysing them. The AMM7 NEMO-ERSEM modelling system has been used to assess impact of climate change on the NE Atlantic in several studies [3, 5, 6], including in the Ocean Acidification assessment included in the last Quality Status Report of OSPAR [7].

#### 1.2 Uncertainty range for fish biomass projections

To address the uncertainty in our NBSS-derived projections of supportable fish biomass, we used the 95% confidence intervals of the underlying CHLa-NBSS relationship from Atkinson et al. [8]. The upper and lower 95% confidence limits of the NBSS slopes were calculated for each grid cell and then used in combination with depth-integrated phytoplankton biomass to estimate the corresponding upper and lower bounds of supportable fish biomass (see methods in main text). These absolute biomass estimates were then converted into percentage changes relative to the 1990-1999 baseline (Figure S3).

#### 1.3 Calculating global averages

Global mean averages were calculated from gridded datasets for projected changes in supportable fish biomass (see methods in main text) and their corresponding confidence intervals (see section above). Grid-cell surface area decreases with increasing latitude, so to account for this we calculated the area-weighted mean from gridded datasets; the value of each grid-cell was weighted by the cosine of its latitude (radians) – a value directly proportional to the grid-cell's surface area.

### 2 Supplementary results

#### 2.1 ERSEM comparison

We compared the phytoplankton and biomass projections from the high-resolution NEMO-ERSEM model against the global CMIP6 Earth System Models used in our main analysis. Phytoplankton projections from NEMO-ERSEM and the CMIP6 models used here broadly agree, resulting in similar projections to future supportable fish biomass in the NE Atlantic (Figure S4). However, NEMO-ERSEM projections include areas of finer-scale increases to supportable fish biomass which are not captured by CMIP6 (Figure S4a, b). Notwithstanding these issues, there is broad model consensus over the overall magnitude of change (Figure S4c).

### 3 Supplementary figures and tables

**Table S1 - Individual sensitivity and adaptive capacity metrics for each country, with corresponding percentile ranks and overall indicator scores.**

Table S1 is attached separately.

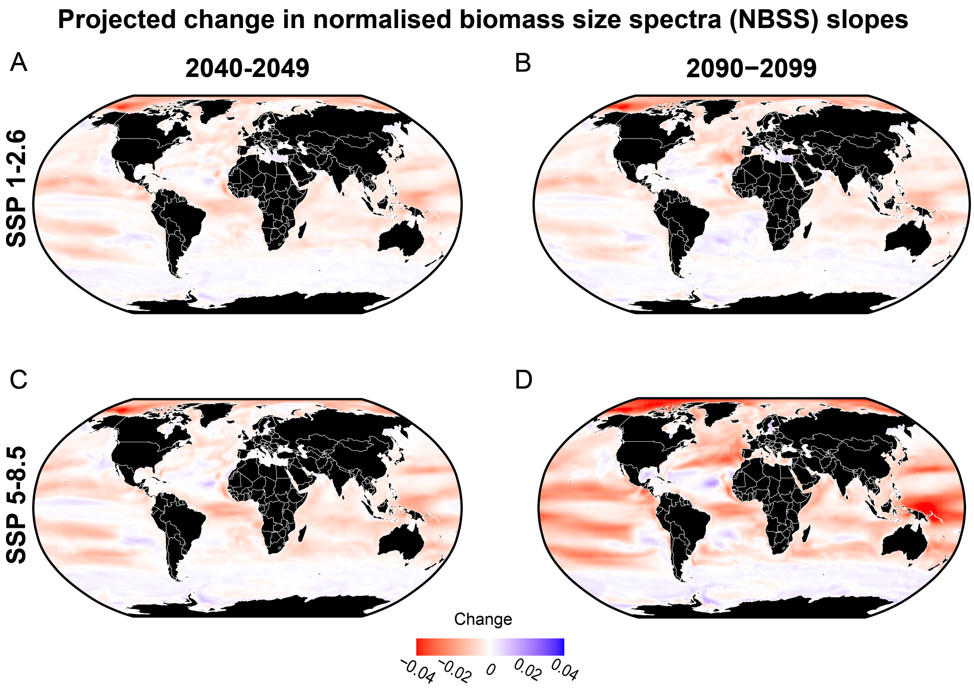


**Figure S1 - Projected absolute changes in normalised biomass size spectra slopes (see methods in main text) relative to 1990-1999.** Absolute changes in normalised biomass size spectra (NBSS) slopes between 1990-1999 and 2040-2049 under **a**). SSP 1-2.6 and **b**). SSP 5-8.5; between 1990-1999 and 2090-2099 under **c**). SSP 1-2.6 and **d**). SSP 5-8.5.

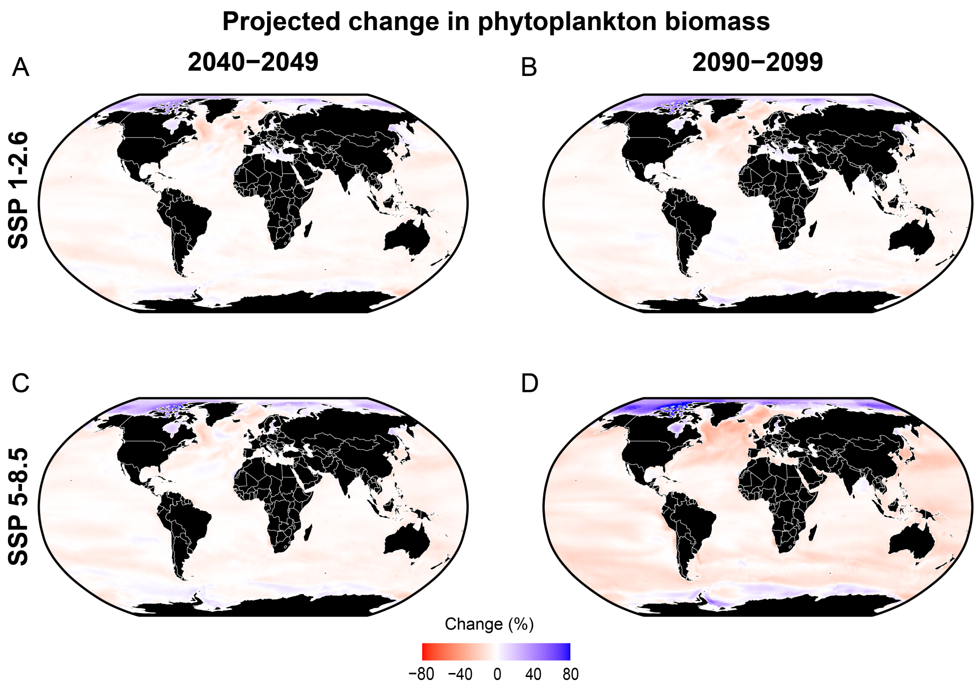


**Figure S2 - Projected percentage changes in depth-integrated phytoplankton biomass from the Earth System Models used in this study (see methods in main text) relative to 1990-1999.** Percentage changes in depth-integrated phytoplankton biomass between 1990-1999 and 2040-2049 under **a**). SSP 1-2.6 and **b**). SSP 5-8.5; between 1990-1999 and 2090-2099 under **c**). SSP 1-2.6 and **d**). SSP 5-8.5.

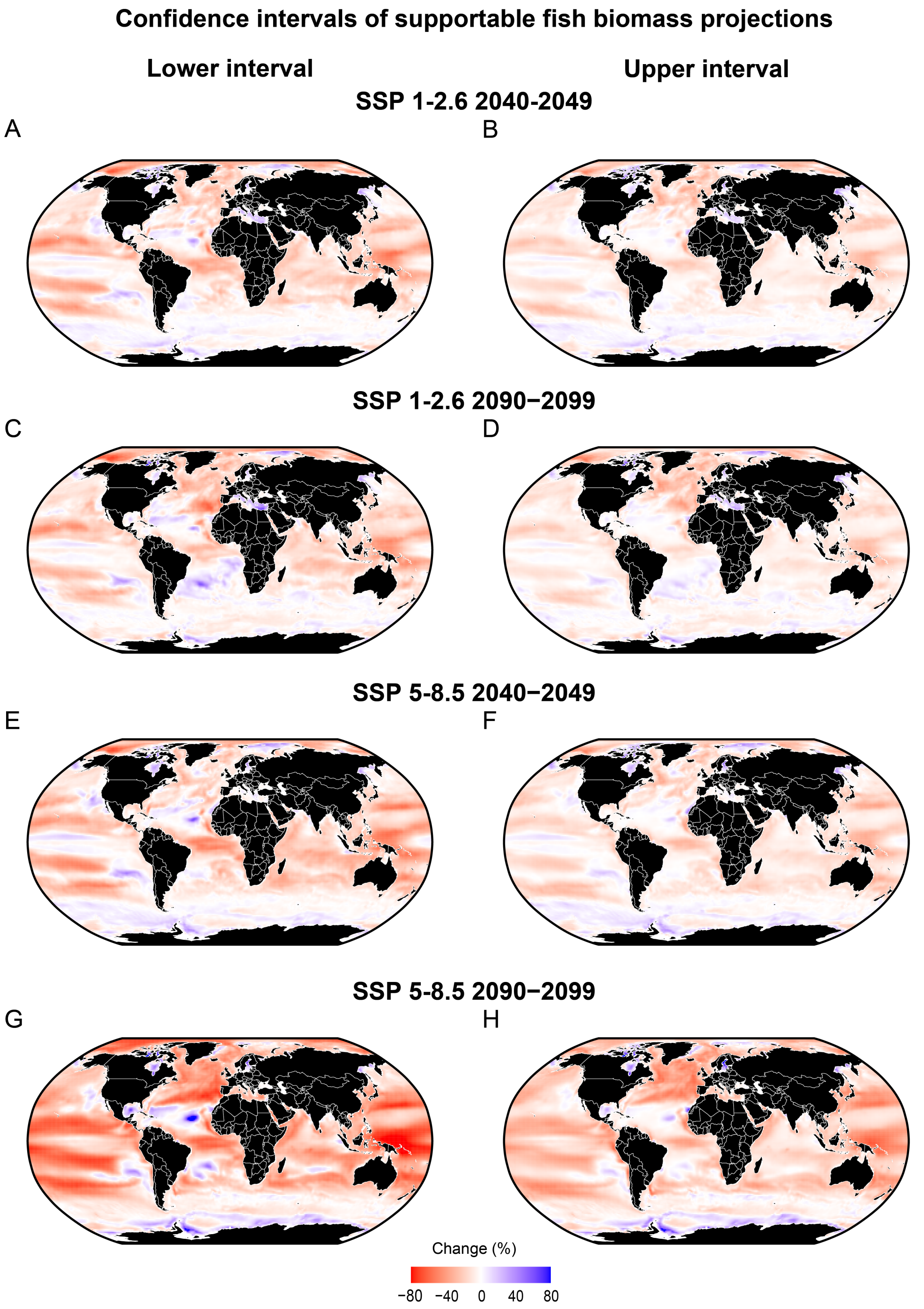


**Figure S3 - 95% confidence intervals of projected changes in supportable fish biomass relative to 1990-1999, based on empirical relationship between Chlorophyll-a and size spectrum slopes presented in Atkinson *et al.*, 2024 (see supplementary methods).** Lower and upper confidence intervals of projected percentage changes in supportable fish biomass relative to 1990-1999 under SSP 1-2.6 (**a**-**d**) and SSP 5-8.5 (**e**-**h**).

**Figure S4 - Hazard and risk scores derived from individual FishMIP models and our NBSS approach.**

Figure S4 is attached separately.

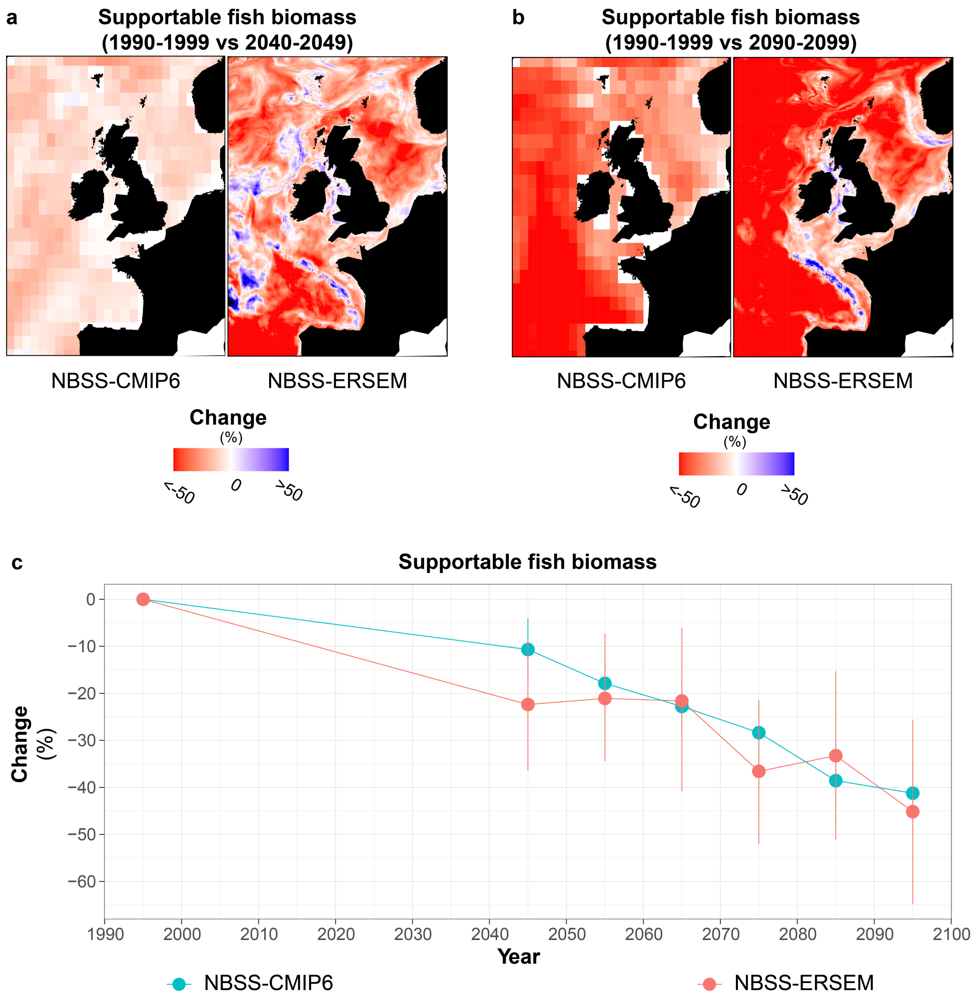


**Figure S5 - Projections of declining fish biomass across the Northeast Atlantic from global scale Earth System Models are supported by high-resolution regional modelling.** Projected changes in supportable fish biomass in the Northeast Atlantic by **a).** 2040-2049 and **b).** 2090-2099. Projections are based on NEMO-ERSEM and CMIP6 Earth System model outputs of phytoplankton changes relative to 1990-1999 under a high emissions scenario (Shared Socioeconomic Pathway 5-8.5 for CMIP6 and Representative Concentration Pathway 8.5 for NEMO-ERSEM), from which our size spectrum slope method was used to estimate changes in fish biomass. **c).** Projected changes in total values over time across the whole map domain (see Supplemetary Methods). Decadal averages were plotted as mid-decade (e.g., 1990-1999 is plotted as 1995).

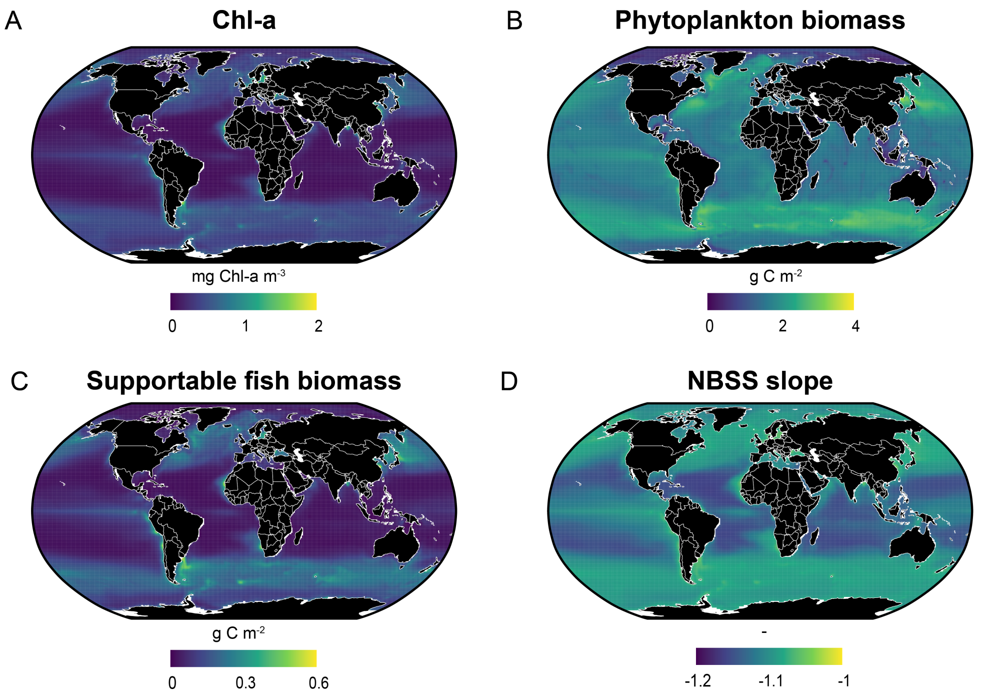


**Figure S6 - Baseline (1990-1990 mean average) values for the variables used to make our NBSS projections of supportable fish biomass (see methods in main text).** Historical (1990-1999 mean average) Earth System Model outputs for **a**). Chlorophyll-a, **b**). phytoplankton biomass, **c**). supportable fish biomass, and **d**). normalised biomass size spectra (NBSS) slopes.

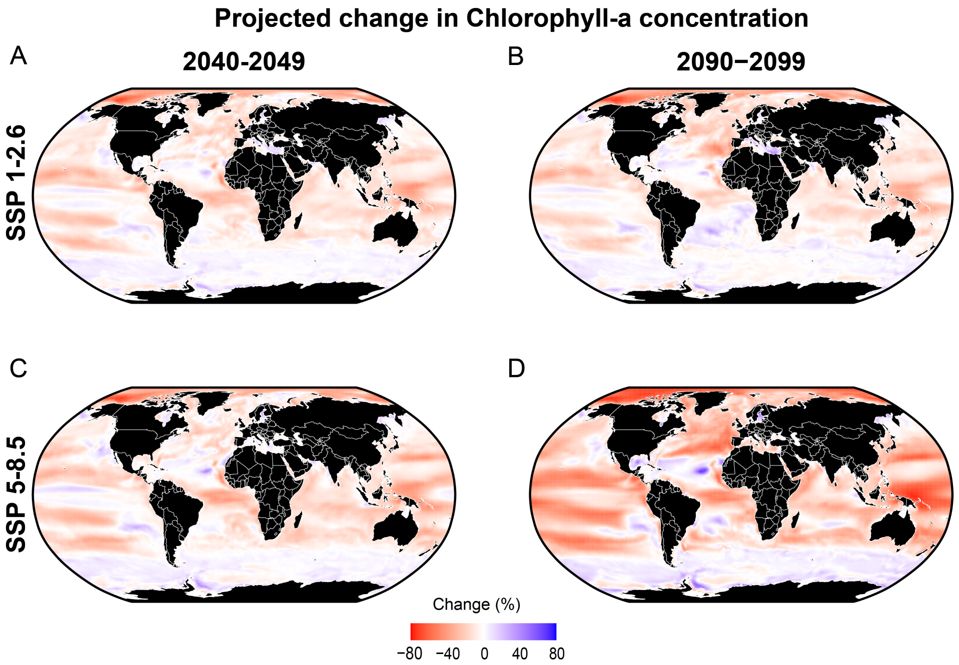


**Figure S7 - Projected percentage changes in Chlorophyll-a concentration from the Earth System Models used in this study (see methods in main text) relative to 1990-1999.** Percentage changes in Chlorophyll-a concentration between 1990-1999 and 2040-2049 under **a**). SSP 1-2.6 and **b**). SSP 5-8.5; between 1990-1999 and 2090-2099 under **c**). SSP 1-2.6 and **d**). SSP 5-8.5.

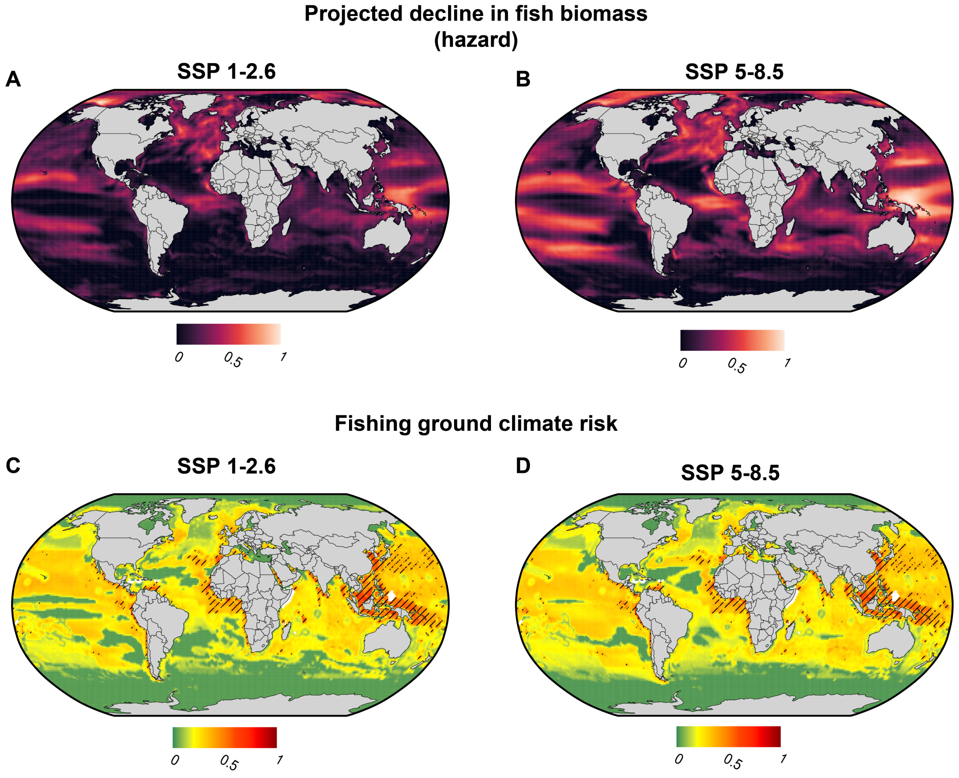


**Figure S8 - The impact of different emissions scenarios on our hazard and risk metrics.** Projected linear trend in supportable fish biomass decline (hazard) between 1990-1999 and 2090-2099 under SSP 1-2.6 (**a**) and SSP 5-8.5 (**b**), derived from size spectra model. Overall fishing ground climate risk under SSP 1-2.6 (**c**) and SSP 5-8.5 (**d**) . Risk values were transformed to enhance visualisation ($y = x^{0.15}$). Hatching (**c** and **d**) indicates regions where both our size spectra projections and the ensemble mean of FishMIP projections (Tittensor *et al.*, 2021) identify the location as high risk ($90^{th}$ percentile). All metrics (**a**-**d**) were rescaled between 0 and 1 for plotting. Missing data are shown in white (see methods in main text).

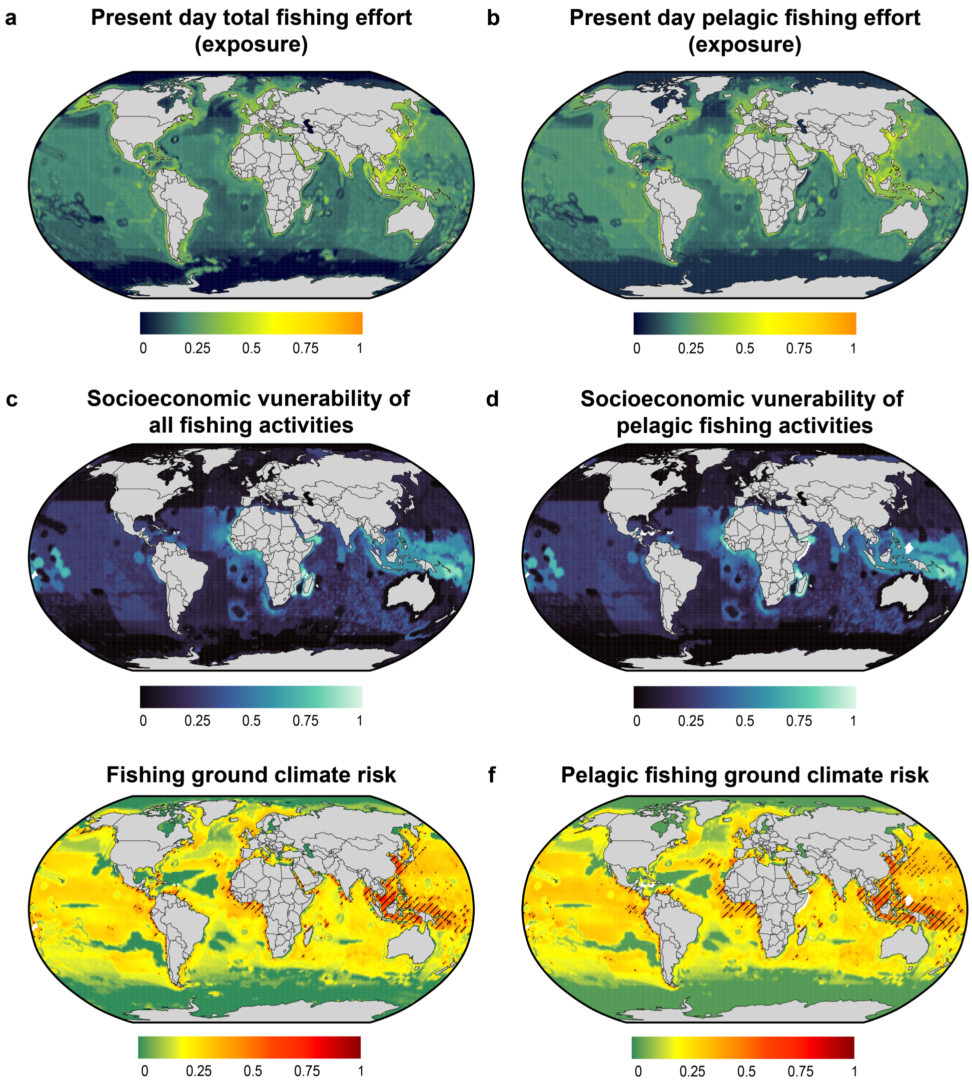


**Figure S9 - Comparing exposure, vulnerability and risk metrics for all fishing activities versus only pelagic fishing activities.** Mean total annual nominal fishing effort between 2014-2017 (exposure) for **a**). all fishing activities and **b**). only fishing activities targetting pelagic fish, processed from Rousseau *et al.* (2024). Exposure values were transformed ($y = x^{0.15}$) to enhance visualisation. Socioeconomic vulnerability of fishing activities weighted by **c**). all fishing activities and **d**). only fishing activities targetting pelagic fish (see methods). Overall fishing ground climate risk calculated with **e**). all fishing activities and **f**). only fishing activities targetting pelagic fish. Risk values were transformed to enhance visualisation ($y = x^{0.15}$). Hatching (**e** and **f**) indicates regions where both our size spectra projections and the ensemble mean of FishMIP projections (Tittensor *et al.*, 2021) identify the location as high risk (90$^{th}$ percentile). All metrics (**a**-**f**) were rescaled between 0 and 1 for plotting. Missing data are shown in white.
